# Novel computational deep learning strategy for neuroprotection identification reveals unique set of nicotine analogs as potential therapeutic compounds against Parkinson’s disease

**DOI:** 10.1101/740050

**Authors:** Felipe Rojas-Rodríguez, Carlos Morantes, Andrés Pinzón, George E. Barreto, Ricardo Cabezas, Leonardo Mariño, Janneth González

**Affiliations:** Departamento de Nutrición y Bioquímica, Pontificia Universidad Javeriana. Bogotá D.C, Colombia; Departamento de Biología, Universidad Nacional de Colombia. Bogotá, Colombia; Instituto de Genética, Universidad Nacional de Colombia, Bogotá, Colombia; Department of Biological Sciences, University of Limerick, Limerick, Ireland; National Center for Biotechnology Information, National Library of Medicine, National Institutes of Health, 8600 Rockville Pike, Bethesda, MD 20894, USA

**Keywords:** Neuroprotection, Nicotine analogs, PI3K/AKT, alpha-7 Nicotine Acetylcholine Receptor (α7 nAChRs), Markov Chain Monte Carlo, Molecular docking, Artificial Neural Network, Parkinson’s disease

## Abstract

Dopaminergic replacement has been used for Parkinson’s Disease (PD) treatment with positive effects on motor symptomatology but with low effects over disease progression and prevention. Different epidemiological studies have shown that nicotine consumption decreases PD prevalence through the activation of neuroprotective mechanisms. Nicotine-induced neuroprotection has been associated with the overstimulation of intracellular signaling pathways (SP) such as Phosphatidyl Inositol 3-kinase/Protein kinase-B (PI3K/AKT) through nicotinic acetylcholine receptors (e.g α7 nAChRs) and the over-expression of the anti-apoptotic gene Bcl-2. Considering its harmful effects (toxicity and dependency), the search for nicotine analogs with decreased secondary effects, but similar neuroprotective activity, remains a promissory field of study. In this work, a computational strategy integrating structural bioinformatics, signaling pathway (SP) manual reconstruction, and deep learning was performed to predict the potential neuroprotective activity of a series of 8 novel nicotine analogs over the behavior of PI3K/AKT. We performed a protein-ligand analysis between nicotine analogs and α7 nAChRs receptor using geometrical conformers, physicochemical characterization of the analogs and developed a manually curated neuroprotective dataset to analyze their potential activity. Additionally, we developed a predictive machine-learning model for neuroprotection in PD through the integration of Markov Chain Monte-Carlo transition matrix for the SP with synthetic training datasets of the physicochemical properties and structural dataset. Our model was able to predict the potential neuroprotective activity of seven new nicotine analogs based on the binomial Bcl-2 response regulated by the activation of PI3K/AKT. We present a new computational strategy to predict the pharmacological neuroprotective potential of nicotine analogs based on SP architecture, using deep learning and structural data. Our theoretical strategy can be further applied to the study new treatments related with SP deregulation and may ultimately offer new opportunities for therapeutic interventions in neurodegenerative diseases.

**Author Summary:** Parkinson’s disease is one of the most prevalent neurodegenerative diseases across population over age 50. Affecting controlled movements and non-motor symptoms, treatments for Parkinson prevention are indispensable to reduce patient’s population in the future. Epidemiological data provide evidence that nicotine have a neuroprotective effect decreasing Parkinson prevalence. By interacting with nicotine receptors in neurons and modulating signaling pathways expressing anti-apoptotic genes nicotine arise as a putative neuroprotective therapy. Nevertheless, toxicity and dependency prevent the use of nicotine as a suitable drug. Nicotine analogs, structurally similar compounds emerge as an alternative for Parkinson preventive treatment. In this sense we developed a quantitative strategy to predict the potential neuroprotective activity of nicotine analogs. Our model is the first approach to predict neuroprotection in the context of Parkinson and signaling pathways using machine learning and computational chemistry.

## INTRODUCTION

Parkinson’s Disease (PD) is the second most common neurodegenerative disorder, mainly affecting population over 60 years of age (1). As a complex and multifactorial disease, PD is caused by several factors such as the accumulation of unfolded proteins, genetic predisposition, mitochondrial dystrophies, epigenetic imbalance and unknown environmental factors that increase the degeneration of dopaminergic neurons in the *substantia nigra* of the brain (2–7). Dopaminergic replacement and related pharmacological treatments are among the most common PD therapeutic strategies oriented to a direct or indirect dopaminergic supply restorage (2, 8). Traditional approaches such as levodopa, carbidopa, and dopamine agonists (apomorphine, amantadine, cabergoline, pergolide, etc), restore dopamine concentration, modulate dopaminergic receptors and stimulate postsynaptic receptors (2). However, these strategies lack long-term efficacy and have been unsuccessful to stop or prevent the progression of the disease (8), and most are directed towards treating PD symptoms such as dyskinesia, motor fluctuations and systemic complications that affect the quality of life of the patients (1,2,9).

Many epidemiological studies have demonstrated an inverse association between cigarette smoking and PD risk (10–12). Pre-clinical studies using *invivo* and *invitro* models have shown that nicotine and nAChR agonists are able to protect nigrostriatal and other neuronal cell populations against damage, suggesting their importance as neuroprotective agents (12–15). nAChRs are ligand-gated ion channels mainly activated by acetylcholine (ACh), and are able respond to other molecules such as nicotine (16). These receptors can be classified into 5 muscle nAChR subtypes (*α*1, *β*1, *γ*/*ε*, *δ*) and 12 neuronal nAChR subtypes (*α*2–10, *β*2–4).

The potential neuroprotective activity of nicotine and nicotine agonists has been linked in part to the activation of pentameric α7 nAChRs, which is highly expressed in dopaminergic neurons and other parts of the brain (14, 17). This, in turn, increases intracellular calcium concentration and activates downstream SP such as extracellular signal-regulated mitogen activated protein kinase (ERK/MAPK), JAK2/STAT3, calmodulin (CAM) and PI3K/AKT. This activation leads in turn to the overexpression of cell survival proteins such as Bcl-x, CREB (cAMP response element-binding) and Bcl-2 (B-cell lymphoma 2) that are related with an increase in neuronal survival, synaptic plasticity and decreased apoptosis (17–19). Moreover, it has been shown that tumor suppressor p53 directly induces Bax transcription, which in turn can overcome the antiapoptotic effects of Bcl-2. These results suggest that p53 regulates the ratio of Bax: Bcl-2 protein levels and therefore influencing the apoptotic fate of a cell in response to stress (20).

Different studies suggest that Bcl-2 is a candidate for the nicotine-mediated neuroprotection in neuronal populations. For example, it has been shown that nicotine induces the phosphorylation of Bcl-2 through either protein kinase C (PKC) and the MAPKs ERK1 and ERK2 or through PI3K-induced phosphorylation of AKT causing the inhibition of apoptosis in neuronal cultures (14,17,18,21–25). Moreover, it has been shown that the overexpression of Bcl-2 protects from neuronal death in mouse (*Mus musculus*) models (26), and the inhibition of this protein results in loss of viability in dopaminergic cells through the induction of caspase-3 (26, 27).

Neuroprotective mechanisms associated with nicotine and nAChRs agonists can prevent the secondary effects of classical therapies such as levodopa-induced dyskinesia, increase dopamine release and improve cytochrome P450 enzymes (CYPs) activity (28, 29). With the advent of nicotine as a potential therapeutic molecule (10–12), toxicity and addictive properties limit its use as a pharmacological agent (30). For these reasons, nicotine analogs retaining neuroprotective activity, but lacking nicotine toxicity and addiction offer an excellent pharmacological alternative in PD (31–35). Previously, we investigated the antioxidant potential of two nicotine analogs against rotenone-induced ROS generation in a PD *in-vitro* model (10). We found that 10 µM of (E)-Nicotinaldehyde O-Cinnamyloxime was able to reduce superoxide anion and hydrogen peroxide production in neuronal SH-SY5Y cells treated with rotenone for 24h (10). This finding is consistent with other studies showing that nicotine analogs decrease superoxide anion generation and oxidative stress in rodent (*Mus musculus* and *Rat ratus*) and primate (*Macaca fascicularis* and *Saimiri sciureus*) animal models (Quik et al., 2012) through the activation of α7 nAChRs, and PI3K/AKT SP (17, 35–38).

Several computational methods such as quantitative structure-activity relation, protein simulation, pharmacophore modeling, and pharmacodynamic models have been used to infer the putative activity of candidate molecules for pharmacological purposes (39). However, in several instances, a systemic approach is required, due to the intrinsic complexity of many diseases such as PD (40). In this aspect, the development of holistic computational methodologies of SP and their modulatory mechanisms can lead to the discovery of new pharmacological targets in complex disorders such as PD (41, 42). In this aspect, the use of ANN (artificial neural networks) have been previously used in a simplified model of the Toll-like receptor signaling pathway, and the PI3K/AKT pathway in the context of cancer (43).

In the present study, we evaluated the potential effectiveness of 8 novel nicotine analogs as possible neuroprotective agents by acting on PI3K/Akt SP. Through the integration of structural and systemic modeling with artificial neural networks (ANN) approaches, it was possible to determine the potential of the nicotine analogs as neuroprotective agents and improve drug scanning efficiency. Even though ANN has been used to study gene interactions (44) or signaling in cancer (43), we use ANN as part of a novel computational strategy aiming to predict novel neuroprotective therapeutic compounds. We present a diagrammatic overview of the interaction algorithm summarizing the presented strategy allowing reproducibility and robustness of the approach (Fig 1).

**Fig 1.**
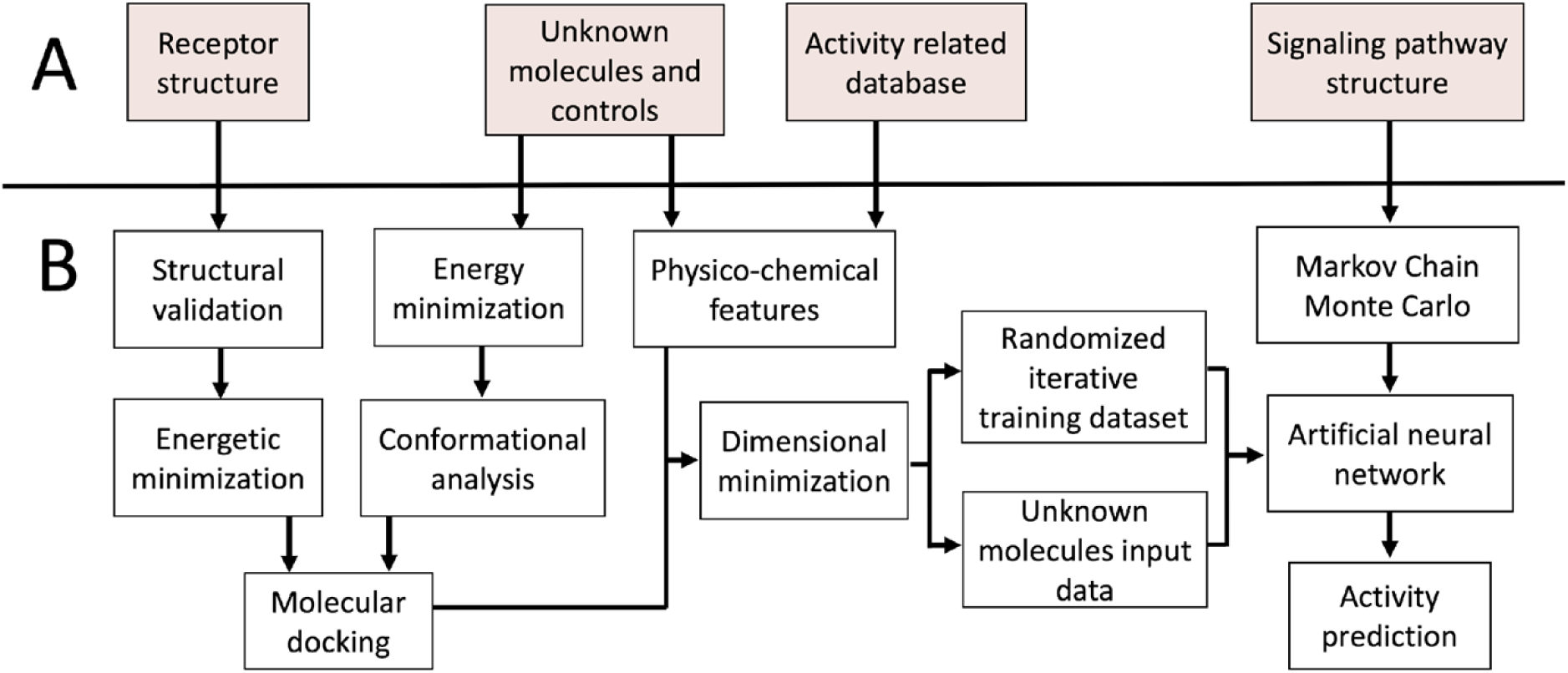
Computational theoretical methodological strategy for ligand prediction over SP architecture. A) Input of biological information B) Computational methods to predict the activity of a series of ligands.

## RESULTS

### Geometry and Structure Analysis

Structures of (3R,5S)-1, methyl-5-(piridine-3-yl) pirrolidine-3-ol (A1), 3-(1,3-dimethyl-4,5-dihidro-1h-pirazole-5-yl) piridine (A2), 3-(3-methyl-4,5-dihidro-1h-pirazole-5-yl) piridine (A3), 3(((2S-4R)-1,4-dimethylpirrolidine-2-yl)) (A4), 3-((2S,4R)-4-(fluoromethyl)-1-methylpirrolidine-2-il)piridine (A5), 3-((2S,4R)-4-methoxi-1-methylpirrolidine-2-yl) piridine (A6), 3-((2S,3S)-1,3-dimethylpirrolidine-2-yl) piridine (A7) and 5-methyl-3-(piridine-3-yl)-4,5-dihidroisoxazole (A8) were used for subsequent methods (Table 1). The neutral molecules optimized at the B3LYP/631G level and conformationally analyzed at the PM6 level showed rotations around C1-C1’bonds and bonds between radical atoms and rings. The minimum energy geometry found was used as an input for the nicotine analogs series and the corresponding energy values (kJ/mol) for the local minimum geometrical conformers were reported in Table 1. The geometrical rotations around the bonds of the rings and the radicals were categorized as true minima of the potential energy of the surface based on the absence of imaginary vibrational frequencies.

**Table 1.**
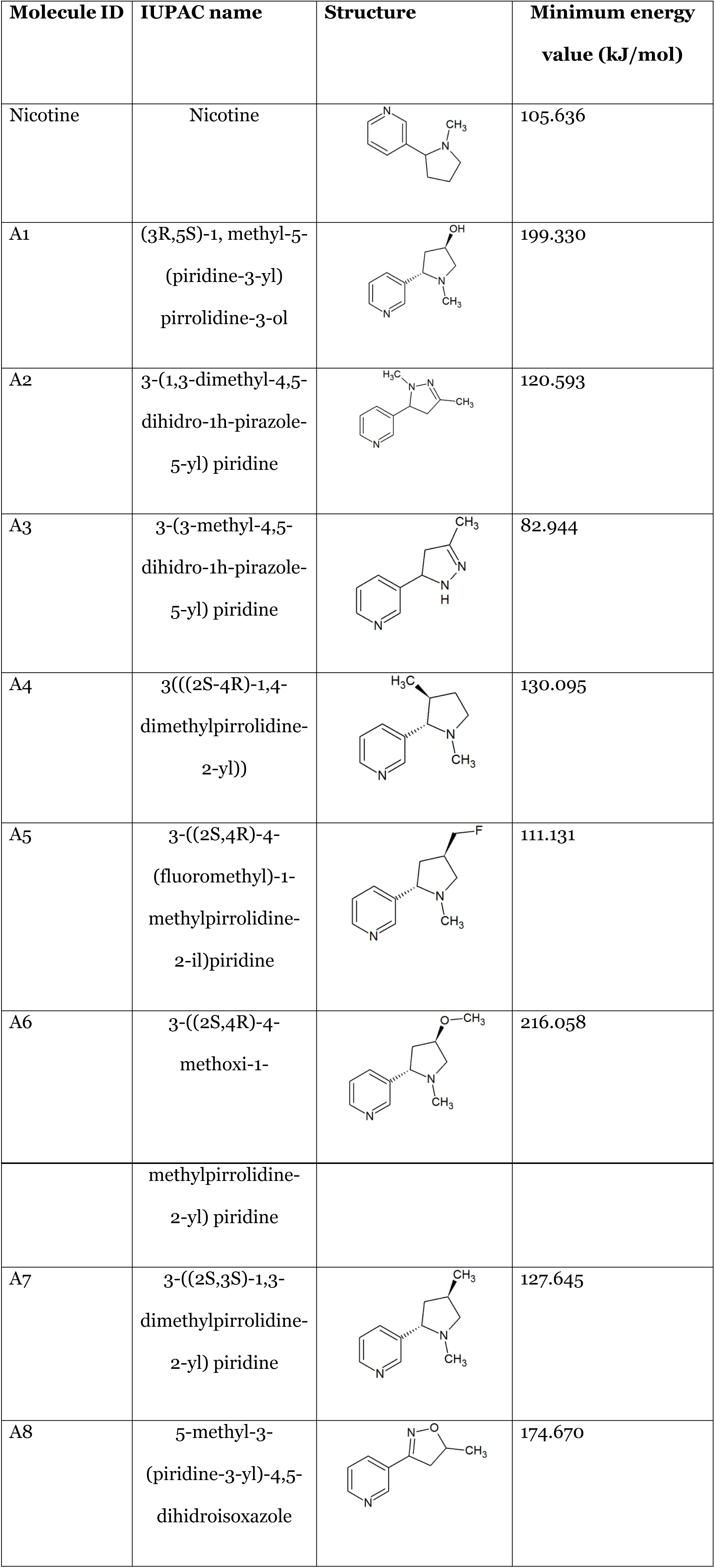
Plane structure and energy (kJ/mol) values of the nicotine analogs. Notice the structural similarity with nicotine due to energy values and the pyridine and pyrrolidine rings present in their structures.

### Nicotine analogs docking and α7-nAChR binding interaction

AChBP ligand-binding structure possesses conserved domain residues forming a narrow hydrophobic pocket, including A (Tyr91), B (Trp145), C (Tyr184 and Tyr191), D (Trp53), E (Leu106, Gln114 and Leu116), whereby A, B, and C are present in a different AChBP ligand-binding subunit than D and E (Fig 2). Following the structural analysis, the nicotine analogs were set to interact with the active pocket within the interaction of the α7 nAChR homo-subunits. Nicotine structure resulted in a geometrical docked conformation in which the pyrrole ring is oriented towards the C loop favoring Van der Waals interactions with amino acids TYR91, TRP145, and TYR184 of the chain A, in addition to LEU106 and TRP53 of the chain B (Table 2).

**Fig 2.**
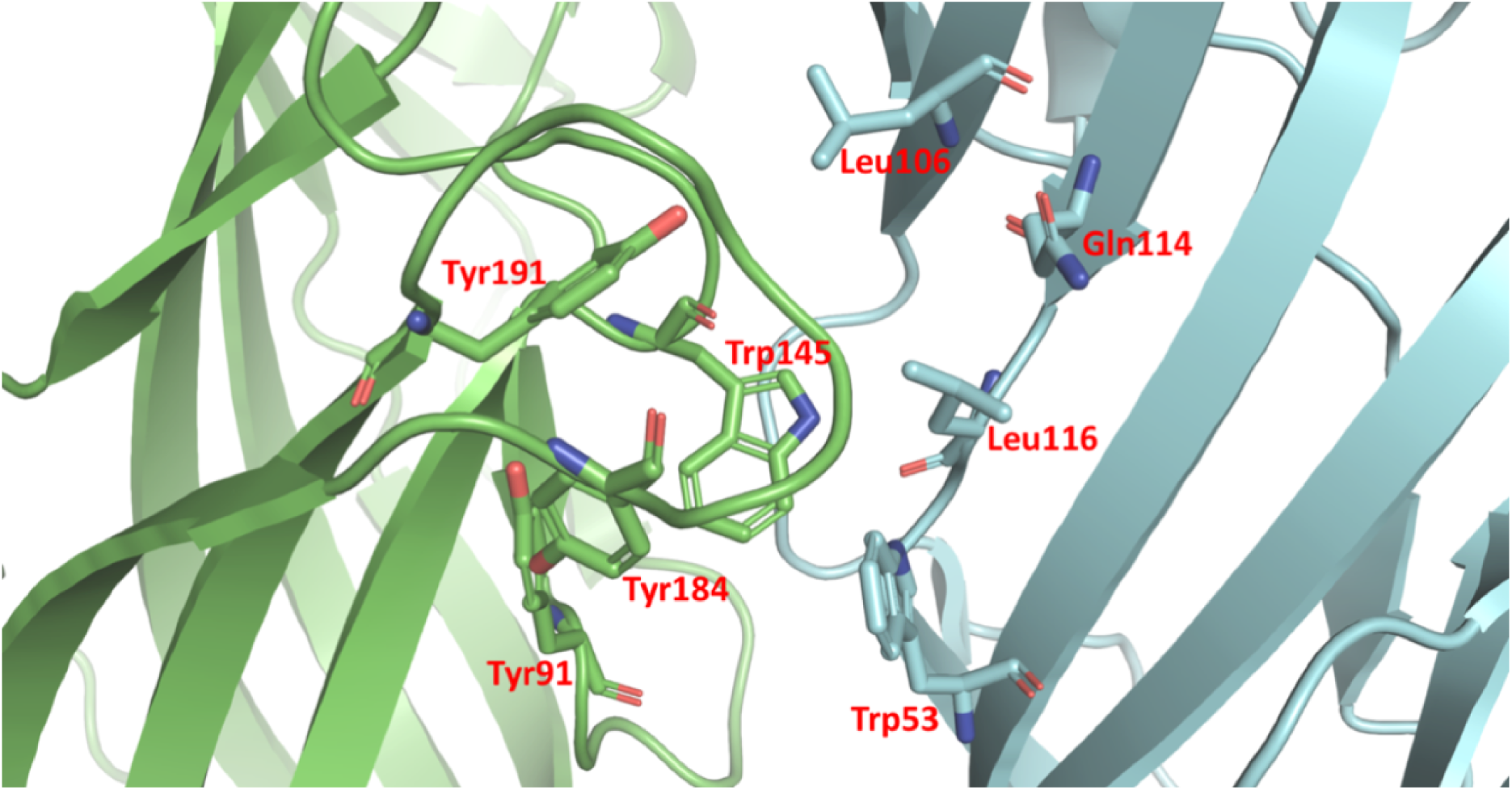
Structural composition of the active site of α7 nAChR. Green and blue represent the two α7 nAChRs subunits composing the active pocket, additional to the localization of the residues composing the active site of the pentameric α7 nAChR.

**Table 2.**
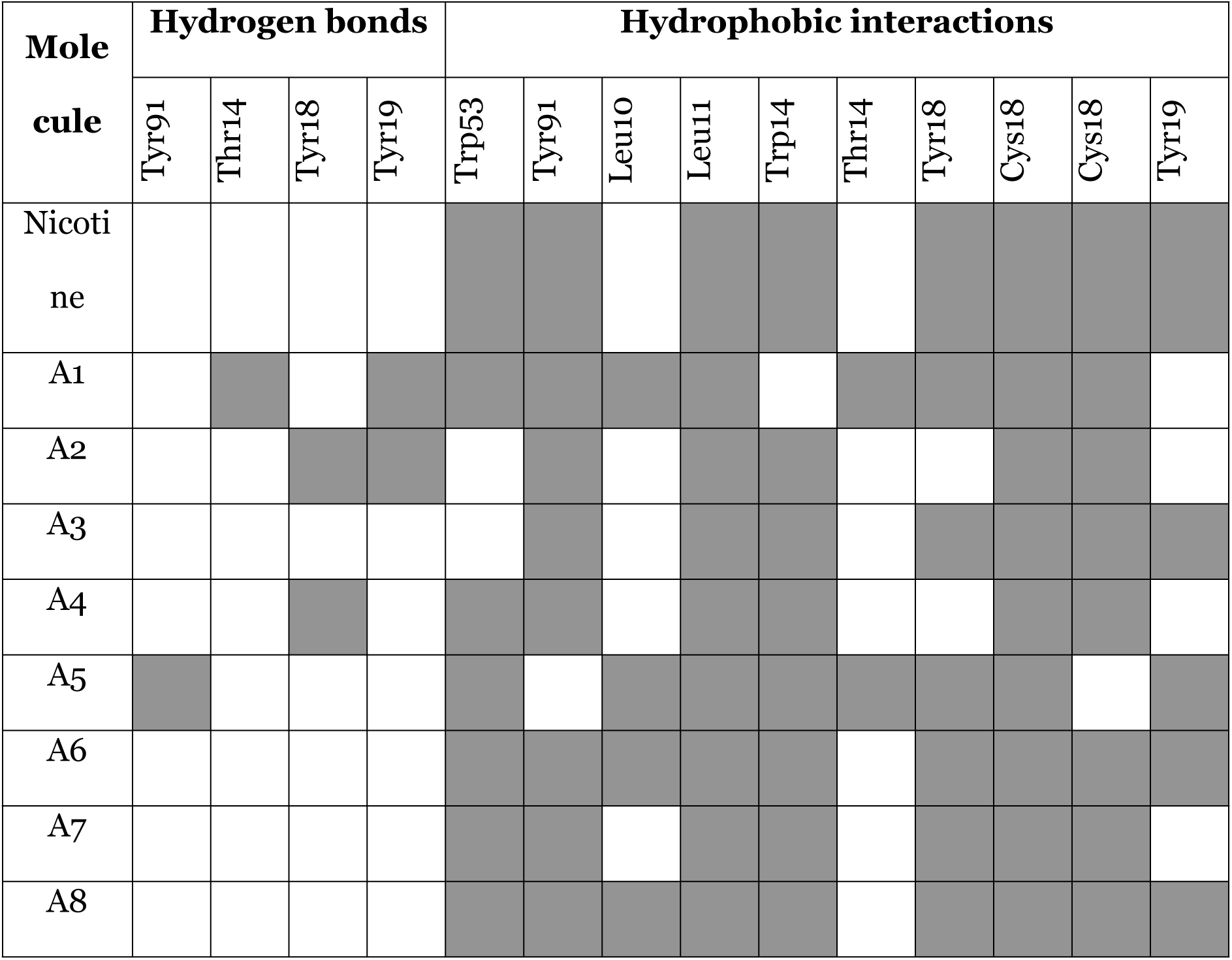
Amino acid residues involved in the receptor interaction with analogs and nicotine. Discrimination between hydrogen bonds and hydrophobic interactions are shown.

Dockings performed with the best ligand conformations were selected and the energy values of interaction are presented in Table 3. The evaluated nicotine analogs, except for the A1 and A3, presented a minor binding energy relative to that of the nicotine molecule itself. Such energetic patterns of interaction between the docked complex allowed a higher affinity in the interaction between analogs and receptor (Table 3). Energetically minimized protein structures interacting with both nicotine and its analogs were used to calculate RMSD in a comparative manner against the docked structure of α7 nAChR interacting with nicotine and wild type protein. Such wild type model was set to be the minimized structure of α7 nAChR without ligand in the active side. The RMSD values for all the protein complexes are presented in Table 3.

**Table 3.** Energy and RMSD values for the molecular docking between analogs and nicotine. For RMSD there is a differentiation between α7 nAChRs interacting with nicotine and α7 nAChRs (wild type) receptor without ligand. For the chimera optimization, the charges for the steepest descent minimization of the ligand were manually added.

| Molecule | Charge of interaction | Energy of interaction (kJ/mol) | RMSD |  |
| --- | --- | --- | --- | --- |
|  |  |  | Wild type | Nicotine |
| Nicotine | +1 | -5.5 | 0,049 | - |
| A1 | +1 | -5.6 | 0,069 | 0,033 |
| A2 | 0 | -5.5 | 0,013 | 0,054 |
| A3 | 0 | -5.6 | 0,039 | 0,075 |
| A4 | +1 | -5.1 | 0,053 | 0,014 |
| A5 | +1 | -4.7 | 0,052 | 0,030 |
| A6 | +1 | -5.5 | 0,047 | 0,016 |
| A7 | +1 | -5.5 | 0,051 | 0,020 |
| A8 | 0 | -5.5 | 0,020 | 0,052 |

### Clustering of similarity

Two manually curated groups of molecules with known agonistic and antagonistic activities were used as input for the cluster. The first group consisted of α7 nAChR agonists SAK3 (45), Nicotine, Acetylcholine, TC-1698 (46), PNU-282987 (Linn, 2016), DMXB (47), and ABT-107 (48). The second group was composed of receptor α7 antagonists and included methyllycaconitine (49), mecamylamine (50), neostigmine (51), anisodamine (52) and bupropion (53). For the second set of molecules we hypothesized a null activation of PI3K/AKT SP, and therefore no neuroprotective activity (54). The optimized conformers for each group were categorized based on general descriptors of physicochemical properties (55, 56). Surprisingly, the dendrogram of similarity was found related with the variables and the neuroprotective activity but with inconclusive scores for similarity (Supplementary Fig S1). Nevertheless, the similarity enriched the experimental data of the ligands using intrinsic structural, physical and chemical values.

### Signaling Pathway reconstruction and predictive model of interaction-response

The reconstructed model of PI3K/AKT SP was able to either induce the activation of the network or produce the inhibition of Bcl-2, as it occurs within the cell (20). Our network is composed of 28 nodes (proteins) that represent relevant components of the pathway (α7 nAChR, PI3K, AKT, CREB, etc), 2 final states (activation/inhibition) and 43 interactions between the nodes (Fig 3). The network was constructed based on nodes with biological relevance associated with the proliferative activity of the network. Using the clustering analysis and the PaDEL-Descriptors previously mentioned, we obtained a set of 1848 variables that characterized physicochemical and pharmacological properties for each one of the molecules. Principal component analysis (PCA, Supplementary Fig S2) and K-mean decomposition (Supplementary Fig S3) were used to reduce the number of variables but maintaining high variance explanation. PCA three-dimensions explained 98.9 % of variance across the original dataset (Supplementary Fig S2) while the K-mean showed a variance explanation of 57.1 % for two-dimensions and the corresponding coordinate values (Supplementary Fig S3).

**Fig 3.**
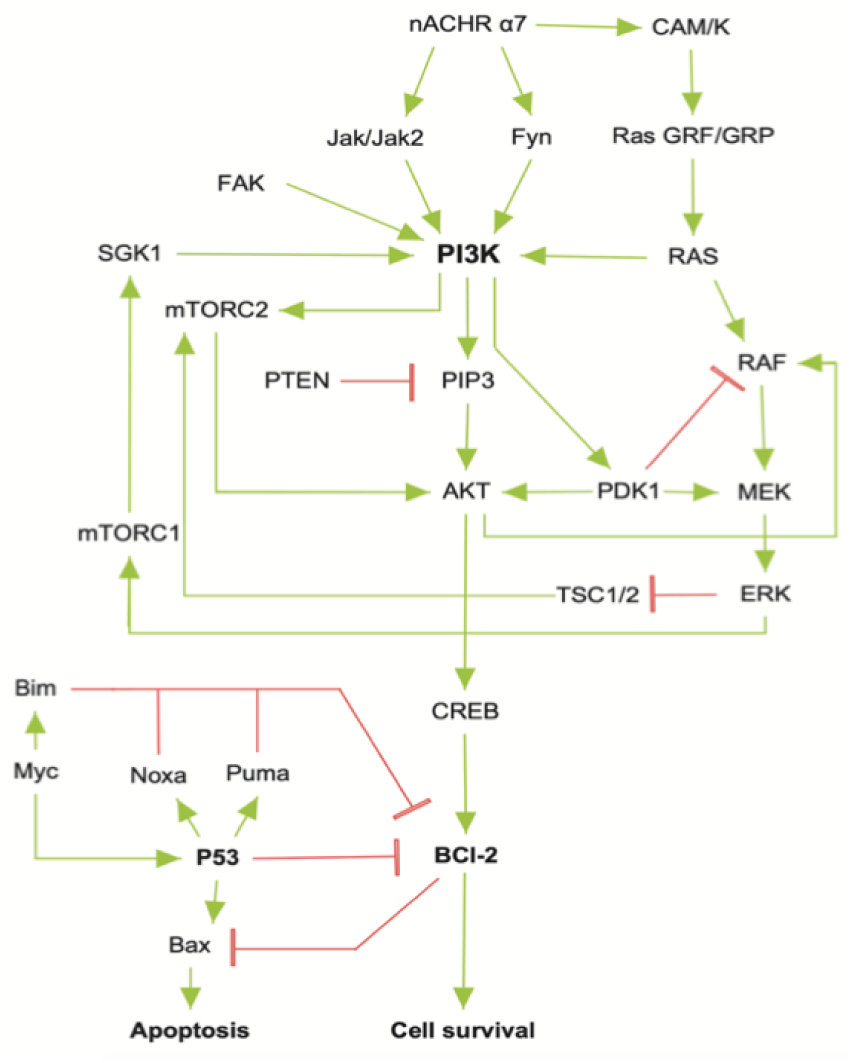
PI3K/AKT Signaling pathway network used to determine the multilayer perceptron architecture. Each node represents the proteins selected for the SP acting over the activity of Bcl-2. Activating and repressing activities were added to model the biological behavior of the SP. P53 was included to mediate a repressive activity over the binomial output.

A consistent MCMC topology was associated with the functional architecture of the reconstructed model representing linear streams in the network (Fig 4). The MCMC model determined the most effective number of hidden layers by iterative topology generation. Convergence was reached with 100.000 iterations resulting in an architecture of 4 hidden layers each one with 5, 1, 4 and 5 nodes, respectively (Fig 5). Based on the architecture of PI3K/AKT represented by the MCMC transition matrix, the multilayer perceptron ANN model was constructed. The topology for the multi-perceptron ANN was particular given that typical architectures for these models tend to reduce the number of nodes per layer across the model (57). We hypothesize that resulting architecture of the model can be associated with the biological topology of the SP and MCMC; nevertheless, additional analysis should be performed.

**Fig 4.**
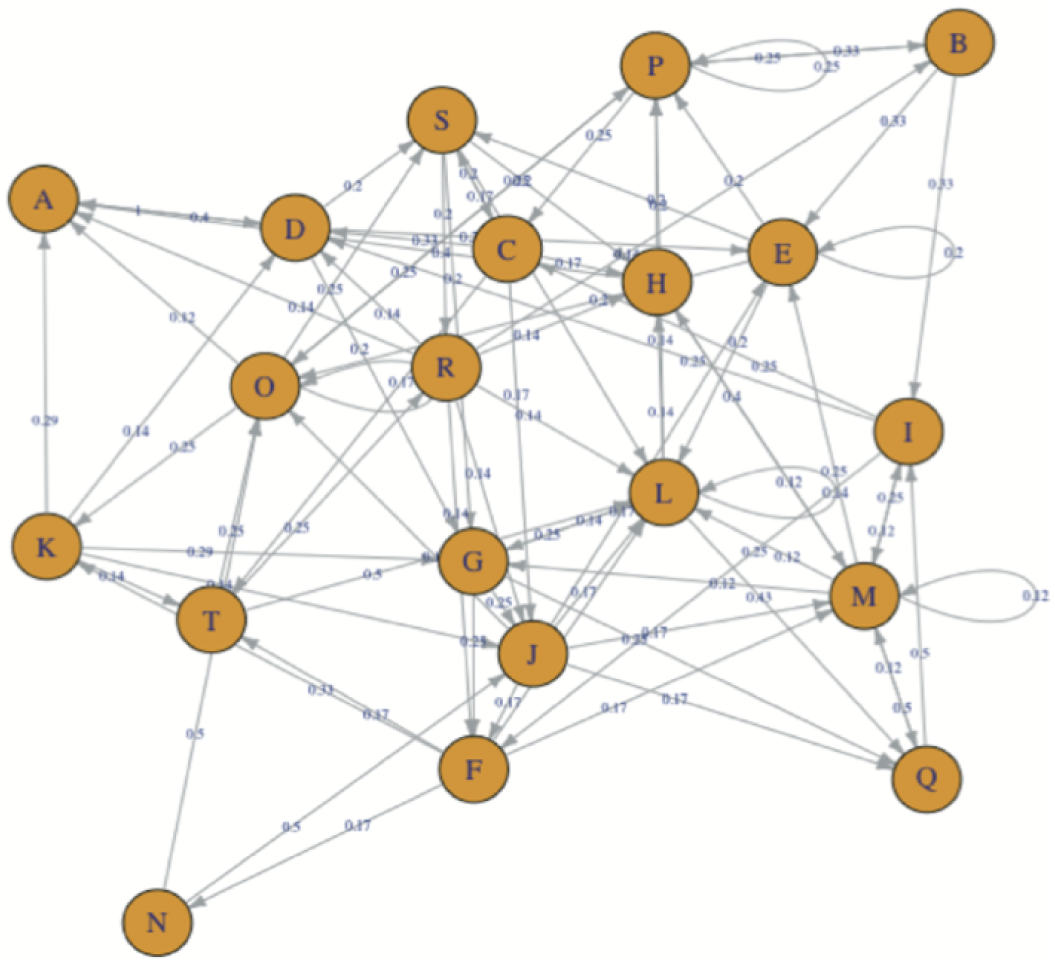
Graphical representation of the probability transitions between nodes in the MCMC model after the 100.000 optimization iterations. Each node represents a state involved in the SP and the edges in the graph establish a relationship between the states.

**Fig 5.**
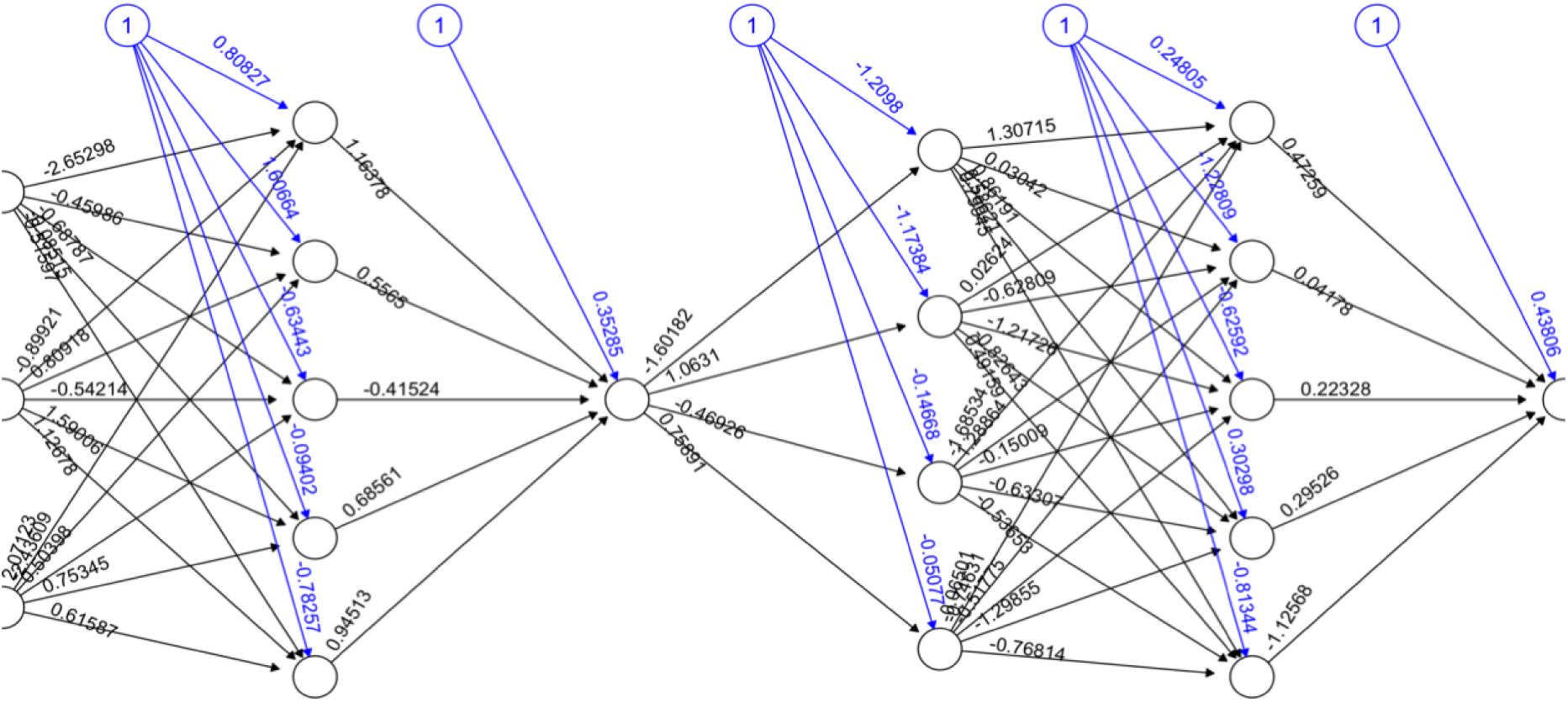
Architecture of algorithm for MCMC-based Artificial Neural Network. The layers and the number of neurons related to each one of the layers is correlated to the architecture of the MCMC optimal topology.

To construct the training datasets for PCA and K-mean associated with each ANN method evaluated, additional PaDEL molecular descriptor variables were added thus increasing sample size. Each method was iterated 1000 times through randomized series and reported values of misclassification (Table 4). Backpropagation with weight tracking coupled with PCA model was capable of generating consistent results across the iterations (Table 5). Our resulting ANN model with for resilient backpropagation with weight backtracking is based on the evolving adaptation rule with a learning process based on error function (58).

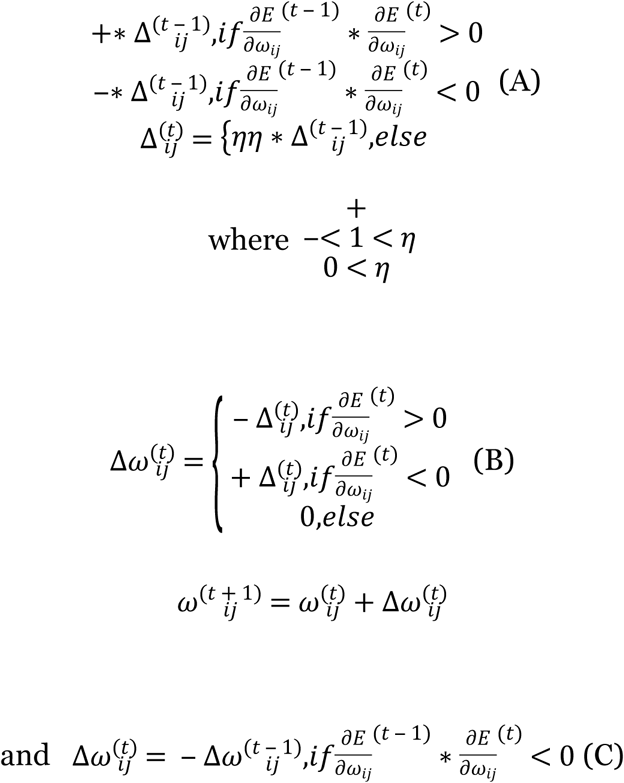

**Table 4.** Error of miss classification for different ANN methods and dimensional minimization approaches. Normalized data is shown (0-1) and the best method is highlighted.

| Learning algorithm | Dimensional reduction method |  |
| --- | --- | --- |
|  | Principal Component Analysis | K-mean |
| Backpropagation | 0.5 | 0.5 |
| Resilient backpropagation with weight backtracking | **<u>0.25</u>** | 0.5 |
| Resilient backpropagation without weight backtracking | 0.375 | 0.25 |
| Smallest absolute derivative | 0.5 | 0.5 |
| Smallest learning rate | 0.5 | 0.5 |

**Table 5.** Predicted values for all the tested molecules. Consistency was evaluated based on the amount of identical values of the prediction over the set of iterations. The binomial output of the model was reported as 1 and 0.

|  | Artificial neural network and dimensional reduction methods integration |  |
| --- | --- | --- |
|  | *Resilient backpropagation with weight backtracking and Principal component analysis* | *Resilient backpropagation without weight backtracking and K-mean* |
| **Consistency** | 100% | 98% |
| **A1** | 1 | 1 |
| **A2** | 1 | 1 |
| **A3** | 1 | 1 |
| **A4** | 1 | 1 |
| **A6** | **1** | **1** |
| **A7** | **1** | **1** |
| **A8** | **1** | **1** |

Briefly, for the adaptation rule each update-value 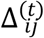 changes depending on the sign for each partial derivative of *ω_ij_* at point *t* (A). If the values are too big then the algorithm skips a local minimum and the function decreases by 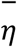 (72). Once 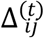 adapts, 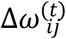 changes according to 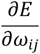 results (B). Positive values of the derivative mean an error increase; therefore, the function is decreased by its 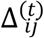. Finally, the exception presented by Riedmiller et al (1993) deals with changes in partial derivative sign (C). In that case the step was too large and missing the minimum, so the previous 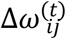 is reverted.

We evaluated the consistency of both the two best models in terms of reproducibility of the predicted results (Table 5). Using this final structure for the multi-perceptron ANN, and the mixture of methods (PCA and Resilient backpropagation with weight backtracking) it was possible to predict the neuroprotective potential of the nicotine analogs. In these aspects, our model predicted that 7 out of 8 analogs had putative neuroprotective activity due to the association of the binomial output of the ANN with the activation of Bcl-2, which is associated with cell survival.

## DISCUSSION

In general, we predicted the potential neuroprotective activity of 7 nicotine analogs in the context of PD therapeutics. To our knowledge, this is the first deep learning neuroprotective computational strategy for ligand molecules activity prediction over α7 nAChR activating PI3K/AKT SP. In general, we presented results supporting the first model capable of reproducing the neuroprotective response of PI3K/AKT induced by α7 nAChRs ligands. By performing conformational analysis, molecular docking, the use of PaDEL variables and geometrical optimization we obtained robust structural conclusive evidence of nicotine analogs activity. Additionally, agonists/antagonist dataset containing experimental data of α7 nAChR receptor interaction and activity allow the replication of *in vitro* experiments. Nonetheless additional data is needed to ensure model robustness and decrease error of misclassification in order to predict non-tested molecules. We now discuss in detail the general aspects of the structural and predictive approaches of the model.

### Conformers’ geometry and ligand-receptor interaction

A considerable amount of research has been done on the pharmaceutical study of nicotine analogs and their molecular structure. However, little assessment has been done about some critical aspects of their conformational variability (15, 59). Recently, the crystallographic structure of the extracellular domain of the nicotinic receptor α7 in complex with epibatidine was reported (60). Although the interactions described for this complex agree with our results, it was found that the residues THR146, CYS187, and CYS186 could also be involved in the protein complex due to their molecular proximity (Fig 2). On the other hand, analog A5 showed a high-affinity score (-4.7) with respect to nicotine (-5.5) (Table 3). This may be due to the ability of pyridine ring to establish hydrogen bonds and the orientation of the atoms in the ligand, particularly with the side chain of the TRP145, similarly to epibatidine. The high affinity of epibatidine has been related to large inter-nitrogen distance previously reported as 4.6 Ǻ (34).

Both nicotine and nicotine analogs showed close behavior and geometry in the 3D conformational structures. For the analogs, variables such as the number of H-bond acceptors and donors were similar with the ones found in nicotine. Two H-bond acceptors and no donor in nicotine are similar to the two H-bond donors of A2, A3, A4 and A7, three for A1, A5, A6 and A8, and only one for A1 and A3. For routable bonds, for A5 and A6 differ from the single bond in nicotine considering the presence of radicals across the pyrrole ring (Table 1). Fluorine radical of A5 and the methyl group attached to the oxygen in the pyrrole ring radical of A6 are rotation-capable bonds relative to the ring plane. Additionally, distinct similarity values in the clustering analysis for analogs A3 and A8 (Supplementary Fig S1) are not reflected on the values of energy affinity or interacting residues. In this case, the clustering is capable of generating a general similarity method based on atom pair descriptors and molecular fingerprints but is inefficient to replicate the docking data. We considered A3 and A8 for the prediction modeling, but further *in vitro* and *in vivo* experiments must be performed to establish if the structural similarities have correlation with the neuroprotective capacity of the tested molecules (Supplementary Fig S1).

Besides the energy affinity, the type and amount of interactions are crucial to determine the role of the docking in the protein function (13). In our model, we found that nicotine interacts by hydrophobic interactions with TRP53, Tyr91, Leu116, Trp145, Tyr184, Cys186, Cys187 and Tyr191 but without hydrogen bonds (Table 2). Compared with the predicted active pocket of the protein, only residues Leu106 and Gln114 were absent in the interaction (61). Nevertheless, hydrophobic interactions with Leu106 were present in the analogs A1, A5, A6, and A8, but no hydrophobic interactions or hydrogen bonds were found with Gln114 (Table 2). The rest of the residues interacting with nicotine were found interacting with the analogs in similar spatial orientations.

None of the analogs showed the same interactions as nicotine, but the energy changes were not higher than 0.4 kJ/mol, except for A5 which has a difference of 0.8 kJ/mol. This change can be associated with electronegative atoms in the ligand, like fluorine and the interaction with Thr146. In this aspect, it is possible that the increase in the energy affinity could be associated with the presence of a hydroxyl radical in the polar sidechain. However, further studies must be performed in order to identify the role of specific residues in the activity of the complex.

For the hydrogen bonds, the presence of two interactions per ligand, A1, and A2 (Table 2) could generate nonspecific responses on activity of the complex, influencing the calcium flow or leading to PI3K/AKT overstimulation. Moreover, fluorine has been associated with secondary effects in humans and, if consumed in higher doses, has been reported to be toxic, thus suggesting that A5 is not a suitable neuroprotective agent (62, 63). The difference in energy affinity, interacting residues and the presence of fluorine lead to the elimination of A5 for extensive analysis because of an increased chance of secondary effects; however, further research must be carried out in order to assess this further. For the RMSD comparison made across all the atoms, after depuration of 25% of the misplaced atoms across the structure, the values for the comparison with the wild type protein and with the protein interacting with nicotine were lower than 0,1. For a comparison between nicotine and the analogs with the wild type the values of RMSD validated our docking result (Table 3). Similar biological response between nicotine, analogs and among analogs can be inferred by the RMSD values of the docked conformations (Table 3).

### Neuroprotective prediction modeling

Only a few studies have employed ANN in the study of signaling pathways (43, 44). For example, (44) developed an ANN of 96 genes with A 3-layered Multi-perceptron with backpropagation learning and sigmoid activation function for the study of biomarkers in children sarcomas (44). Moreover (43) used a simplified neural network to integrate some of the environmental and molecular characteristics of cancer progression. This network was comprised by two microenvironmental input nodes (growth factor and death signal), and two phenotype output nodes (pro-growth and pro-death, (43). However, to our knowledge, this is the first study that combines structural data, docking simulations and ANN in the modulation of a SP.

In our model, it was possible to extract the values related to the putative neuroprotective activity of the analogs through the generation of a suitable dataset using the physicochemical and docking data as reference. Nevertheless, inconclusive relations between ligand structure and complex interaction with the neuroprotective response occurred in the structural clustering. For example, the similarity values across the dendrogram generated an aggrupation node consistent with all the neuroprotective ligands (Supplementary Fig S1). Nevertheless, the ligands without a neuroprotective association did not generate a suitable group but a disperse distribution across the dendrogram. In this sense, the FOF method implemented in the clustering analysis was unable to discriminate among biological effects (proliferation and antiapoptotic) because of the strong aggrupation occurred only for neuroprotection positive molecules (Supplementary Figure S1). Moreover, we discarded acetylcholine, the natural ligand of the receptor and a ubiquitous neurotransmitter in the brain because it was the most divergent ligand of the studied group; however no neuroprotective function has been associated with this molecule (64). Inclusion of synthetic molecules such as SAK3, PNU-282987 and allosteric modulators improves the conclusions considering that they have been previously reported to induce neuroprotection through the modulation of PI3K/AKT/Bcl-2 (14).

By including a higher set of variables, we observed that in the PCA and K-mean decomposition analysis, the analogues values tended to cluster near to nicotine (Supplementary Figs 2 & 3). Importantly, comparing the clustering and dimensional minimization results allowed us to improve the set of chemical variants and the derived synthetic data for further analysis. PCA and K-mean showed a similar behavior for the cluster mixed with other molecules but significant differences in variance representation, being 98.9% for PCA and 57.1% for K-mean (Supplementary Fig 2 & 3). The absence of strong groups in the PCA and K-mean analysis showed that synthetic data was not conclusive to predict the biological activity by themselves. With this in mind, the ANN model was essential to discriminate between the groups and underline fundamental properties of the analogs based on the SP structure.

Using the synthetic values of PCA and K-mean it was necessary to determine a proper SP model to generate reliable predictions. SP are dynamic systems in which crosstalk with neighbor paths is essential to modulate signal intensity (65). The addition of p53 negative feedback increased the robustness of the model by modifying Bcl-2 binomial output and information flow across the model (Fig 3). This negative regulation supports the control of the system and structure, leading to a suitable MCMC path (Fig 4). Considering that the matrix of transition represents the structure of the network and that the probabilities of transition were iterated until stability was reached, the biological regulation of PI3K/AKT by neighbor SP leads to robust computational inferences (66). Additionally, our network has biological relevance integrating the receptor to the PI3K/AKT network showed the biological relevance of ionotropic channels such as α7 nAChR for PI3K/AKT activation without explicitly acknowledging the presence of JAK (14).

Based on the SP diagram and the iterative conjugation of the MCMC transition matrix, we built up the non-canonical multi perceptron hidden layer structure (Fig 5). The presented ANN resulted as a biological-based discrimination of architectures through iterations using MCMC matrix. Similar approaches have been used before to prevent the overfitting of the model. For example, the drop-out technique allows to drop units from the neural network during training hence preventing an excessive coadaptation and overfitting of the network (12). The iterative process was set to converge at the minimal topology with significant predictive capacity of the ANN model. The use of MCMC allowed to reduce the number of layers and nodes by establishing blocks between the input of information and the classification output (67). The hidden layers were locked according to the MCMC product (Fig 4), ensuring that the subsequent topology represented the biological information contained in the SP. In this aspect, the synthetic iterative randomized data sets integrated with dimensional minimization and ANN, allowed the determination of the best predictive method reducing error and ensuring consistent predictions during training.

Using PCA and K-mean is an effective way to effectively decrease the sample variability to a low number of synthetic values (68). As mentioned before, PCA and K-mean methods were capable of reproducing the results of the clustering by reducing the number of variables from 1848 variables to a small dataset. The values used correspond to the 3 main coordinates of the PCA and K-mean methods for each one of the molecules, optimizing time and performance of the method and reducing the over fitting of the ANN model. Through variable reduction, we increased the sensibility and reliability of the model to predict the binomial neuroprotective signal output. Additionally, it was possible to decrease time consumption, allowing the chance of model reproducibility.

In the case of the ANN, there has been an important amount of development in algorithms of learning and error (69). Regarding the methods tested to train and implement our model, we found identical split differences in misclassification for smallest absolute derivative, backpropagation and smallest learning rate with PCA and K-mean, and resilient backpropagation with weight backtracking with K-mean (Table 4). We found an improvement from both resilient backpropagation with weight backtracking with PCA, and resilient backpropagation without weight backtracking with K-mean reducing misclassification to 25%. However, resilient backpropagation without weight backtracking with K-mean was unable to predict a consistent output for the analogs, meaning an output change through the iterations (Table 5). The prediction was consistent for positive neuroprotective output in 98% of the iterations but the remaining 2% was negative in the binomial set of the network. The method of Resilient backpropagation with weight backtracking with PCA resulted as the best combination for neuroprotection prediction, because 100% of the predictions were consistent along the iterations (Table 5). In this case, the resulting model was based on the implementation of an adaptation rule with a learning based on error values purposed by (58). By predicting the binomial neuroprotective output and integrate them with the structural conformational and docking analysis, it was possible to determine that 7 out of 8 analogs are potential neuroprotective agents (Table 5), consistent with their previously reported activity in neuroprotection (10,48–53).

Although the model corrects the inner functions for noise in the data, further testing is essential to determine the true potential of nicotine analogs and antagonist for PD therapy. In this regard, the use of *in-vitro* and *in-vivo* data, in either dopaminergic neuronal cell lines, cerebral organoids or animal models, can contribute to improvement of the model and its predictive capacity. Even so, our model was in accordance with previously publish experimental data (10,48–53) proving to be robust enough as testing method for further research. Nonetheless, additional training data is necessary to improve predictability of the ANN. Furthermore, this strategy can be used to predict the secondary effects of nicotine analogs by dataset modification with addiction/reward molecules like nicotine and methyllycaconitine (49). In this aspect, we suggest that the addition of a bigger dataset of antagonist and agonist molecules could provide a more robust training dataset leading to a reduced error of misclassification. Our strategy (Figure 1) ensures the biological discrimination of the prediction with the architecture of the ANN supported by the SP MCMC. Worth noticing, SP are dynamic systems with continuous variables and therefore, the intermediate states are essential for its appropriate behavior prediction (65). This intermediate state of the SP behavior can be modeled using partial differential equations but the integration with ANN model in SP is still challenging. Finally, further analysis like protein dynamics must be implemented to describe in detail the spatial responses of the receptor to new drug-like molecules. However, we propose that the integration of dynamic models with this novel conceptual strategy could be applied to predict alterations in SP networks in the context of potential neuroprotective mechanisms.

In summary, this theoretical strategy is a novel quantitative approach that could be used to predict the interaction of new molecules that affect PI3K/AKT signaling either on PD or in other pathologies associated with impaired signaling pathways. The algorithms of Resilient backpropagation with weight backtracking with PCA showed to be the best combination in our model to validate the neuroprotective activity of the studied molecules. Additionally, enabling computational testing of nicotine analogs pharmacological properties beyond PD therapeutics is important considering that agonists of α7 nAChR have been reported as promissory candidates for the treatment of Levodopa-induced dyskinesia. Nevertheless, further studies must be performed to establish the full potential of our neuroprotective model and its applications in neurodegenerative research.

## MATERIALS AND METHODS

### Conformational analysis and protein structure preparation

8 novel nicotine analogs (Table 1) were used in the present study: (3R,5S)-1, methyl-5-(piridine-3-yl) pirrolidine-3-ol (A1), 3-(1,3-dimethyl-4,5-dihidro-1h-pirazole-5-yl) piridine (A2), 3-(3-methyl-4,5-dihidro-1h-pirazole-5-yl) piridine (A3), 3(((2S-4R)-1,4-dimethylpirrolidine-2-yl)) (A4), 3-((2S,4R)-4-(fluoromethyl)-1-methylpirrolidine-2-il)piridine (A5), 3-((2S,4R)-4-methoxi-1-methylpirrolidine-2-yl) piridine (A6), 3-((2S,3S)-1,3-dimethylpirrolidine-2-yl) piridine (A7) and 5-methyl-3-(piridine-3-yl)-4,5-dihidroisoxazole (A8). Molecular structures were sketched on Avogadro using MMFF94s forcefield, correcting atom type and chirality (Hanwell, et al., 2012). Calculations were carried out with Gaussian 09 (Frisch et al., 2016), using B3LYP level of theory and cc-PVDZ basis set. In order to find the minimum energetic conformation, the structures were optimized at the DFT B3LYP/6-31G level. Conformational analysis was carried out on minimized rotating bonds between pyrrole, derivate rings, and pyridine rings, including the bonds of all the radicals of the pyrrole ring for each ligand. The maximum number of conformers for each molecule was set to 30, with the 10 lowest energy conformations used for the docking simulations.

The crystal structure of α7 nAChR in complex with lobeline was obtained from the RCSB Protein Data Bank (PDB ID: 5AFJ) (61). AutoDockTools (70) was used to assign polar hydrogens and add Gasteiger charges. The geometry of the receptor was optimized using the MM2 molecular mechanics force field. The neutral and ionized states of aliphatic amine and carboxylic acid groups of compounds to be docked were protonated and deprotonated separately.

### Molecular Docking

To determine the interaction between the studied molecules and the receptor, docking simulations were performed with AutoDock4 (version 4.2) (70). The active site of the pentameric structure of α7-nAChR was defined as the interfaces between subunits that were within 12 Å from the geometric centroid of the ligand (61). Within the 3 domains of the protein (extracellular, intracellular and transmembrane), the pocket is located in the extracellular side of the receptor with residues from loops A-C of the principal subunit and loops D-E of the complementary subunit. Default settings for small molecule-protein docking were used throughout the simulations. Worth noticing, α7 nAChR has 5 active sites located at the subunit interactions of the protein complex. Considering that homopentamer protein has 5 identical active sites, we modeled the interaction within one of the pockets.

Docked conformations were clustered according to the interacting energy combined with geometrical matching quality. The complexes with the best score were taken as the lead conformation for each compound. The correlation between key interactions obtained from the computational simulations and the interactions reported by the crystallographic structure of α7 nAChR in complex with lobeline was further explored. To further analyze the differences between the interaction of the analogs and the receptor, we calculated the RMSD values between analogs, nicotine and the receptor. RMSD was calculated using 85% of the atoms resulted after 3 outlier rejection cycles. The energetically minimized docked structures of the α7 nAChR receptor were made with Chimera (71), using 10000 steepest descent steps, 0.001 Ǻ steepest descent steps size, 10 conjugate gradient steps, 0.001 Ǻ conjugate gradient steps size, update per 10 intervals, gasteiger method for charge and Amber ff14sb algorithms for residues.

### Physicochemical clustering and dimensional decomposition

To determine the features for each molecule and their structural similarity, physicochemical property predictions, classification and clustering of structures were performed on ChemmineR, R package (55).

Structural similarities between analogs were compared to a manually curated dataset of 7 agonists and 5 antagonists of α7 nAChR with reported neuroprotective activity (45–53,72). This method allowed us to increase the size of the available data, robustness and reproducibility. In the dataset, agonists of α7 nAChR were associated with the induction of a neuroprotective pathway, either expressed through the induction of cell proliferation or apoptotic prevention. Antagonists were set to block either the proliferative activity by PI3K/AKT of the signaling pathway or the associated Ca^2+^ mobilization of the receptor (52,53,73,74).

Clustering analysis was based on molecular descriptors such as molecular formula, molecular weight, atom frequency and functional groups. To determine the optimal cluster, a matrix of molecular descriptors for all molecules was used, categorizing the ligands according to the structural, physical and chemical variables. To increase the robustness of the clustering model, additional physicochemical data from *de-novo* featurization was included (56). To ensure a proper physicochemical featurization and increase the number of variables available for further analysis, data from *de-novo* characterization using PaDEL-Descriptors (56) was also included. Dimensionality reduction of the dataset was done by principal component analysis (PCA) and k-mean decomposition. The mentioned approaches were used to identify the multiparametric data capable to describe the agonistic and antagonistic function and the relation between the activity and structure of the ligands in the dataset.

### Signal reconstruction

To establish a reliable topology of the SP related with the neuroprotective capacity of nicotine, a manually reconstructed network for the PI3K/AKT SP was developed using information from KEGG (75–77) and PantherDB (78). To ensure the biological coherence of the network, our model was enriched with additional protein-protein interactions that are present in the PI3K/AKT/mTOR interactive pathway (79). Finally, this PI3K/AKT SP was integrated with a manually curated model of p53 to generate a negative feedback over Bcl-2. In this aspect, the model included 6 specific proteins (Myc, Bim, P53, Bax, Noxa, Puma) associated with the partial inhibition of the Bcl-2, which emulates the SP biological behavior (66).

### Activity prediction and interaction-response model

In order to elucidate the quantitative relationship between α7 nAChR receptor and the activation of Bcl-2, a Markov Chain Monte Carlo (MCMC) model for the whole network was implemented. This model was based on the reconstructed PI3K/AKT SP network and was generated using the R package *MCMCpack* (80). A matrix of transitional stages predicting the most suitable path across the nodes was associated with the end stages of the MCMC that represented Bcl-2 expression as a binomial logical argument (Table 5). The construction of the artificial neural network model (ANN) was performed with the *NeuralNet* package in R (81). To determine the canonical architecture for the ANN we used an optimization algorithm using 100.000 iterations using the MCMC transition matrix as an input. A stable multi-perceptron structure was set to have a minimal number of hidden layers with the capacity of representing the MCMC transition matrix, as shown below:

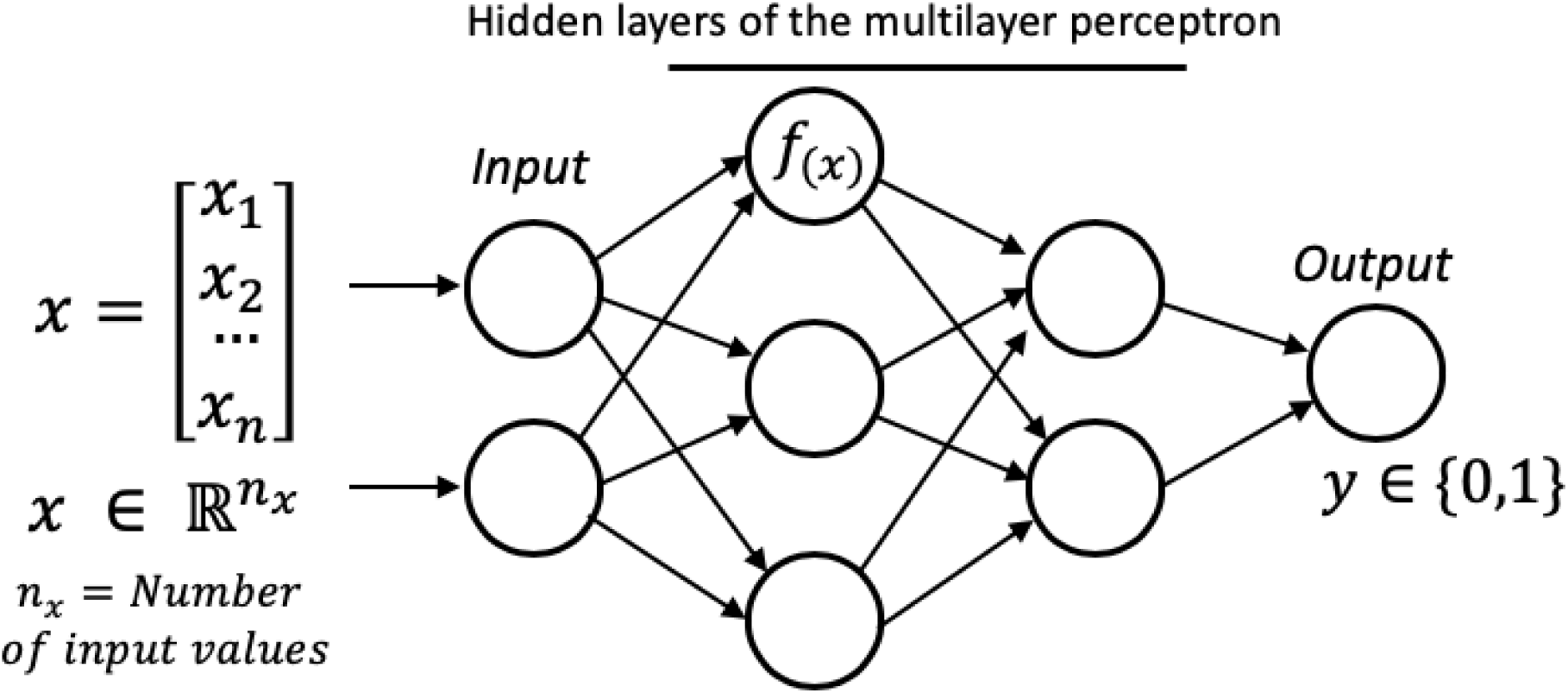

Briefly, a matrix *x* is used as input in the first layer of the multiperceptron. Each node of the hidden layer computes *f_(x)_* to finally generate a binomial *y* ouput.

The dimensional reduction methods were used to train the ANN using the coordinates in the PCA and K-mean. One thousand randomized training datasets were selected and associated with the binomial activity of the ligand dataset. To optimize the predictive capability of the model using the training datasets, four ANN algorithms were compared: Backpropagation with a learning rate of 0.001, Resilient backpropagation with weight backtracking, Resilient backpropagation without weight backtracking, smallest absolute derivative model and smallest learning rate. Additionally, for each possible combination of dimensional minimizations and ANN methods, the random data subsets used as training were iteratively generated, after excluding the testing values. To ensure the robustness of the prediction, values for misclassification error were obtained for each training and testing combination. In this aspect, the best model acquired was set to minimize the error to reduce the amount of false-positive predictions.

## ACKNOWLEDGMENTS

This work was supported by the Pontificia Universidad Javeriana, Bogotá, Colombia, and Colciencias ID 7740 and 7714 to JG. We would like to thank Alix Loaiza for providing nicotine analogs and the DIT support team at Pontificia Universidad Javeriana for the troubleshooting the server related issues.

## Declarations

### Ethics approval and consent to participate

“Not applicable”

### Consent for publication

“Not applicable”

### Availability of data and material

The datasets used and/or analyzed during the current study are available from the corresponding author on request.

### Authors’ contributions

FR-R developed the structural analysis for the ligands with the receptor, the manual reconstruction of the signaling pathway and the dataset for the agonist and antagonist ligands related to the receptor including the physicochemical featurization. CM developed the MCMC decomposition of the PI3K/AKT signaling pathway. CM and FR-R developed the iterative process for the architecture for the ANN and the corresponding training datasets. RC contribute for the writing and biological analysis of the results. AP and JG were major contributors of the biological significance of the results and improved the methods and applications along the development of the project. GEB and LM contributed with the biological impact of the computational framework and applications. All authors contributed to analyzing the ANN data, manuscript writing and approved the final manuscript.

## SUPPLEMENTARY FIGURES

**SUPPLEMENTARY Fig 1.**
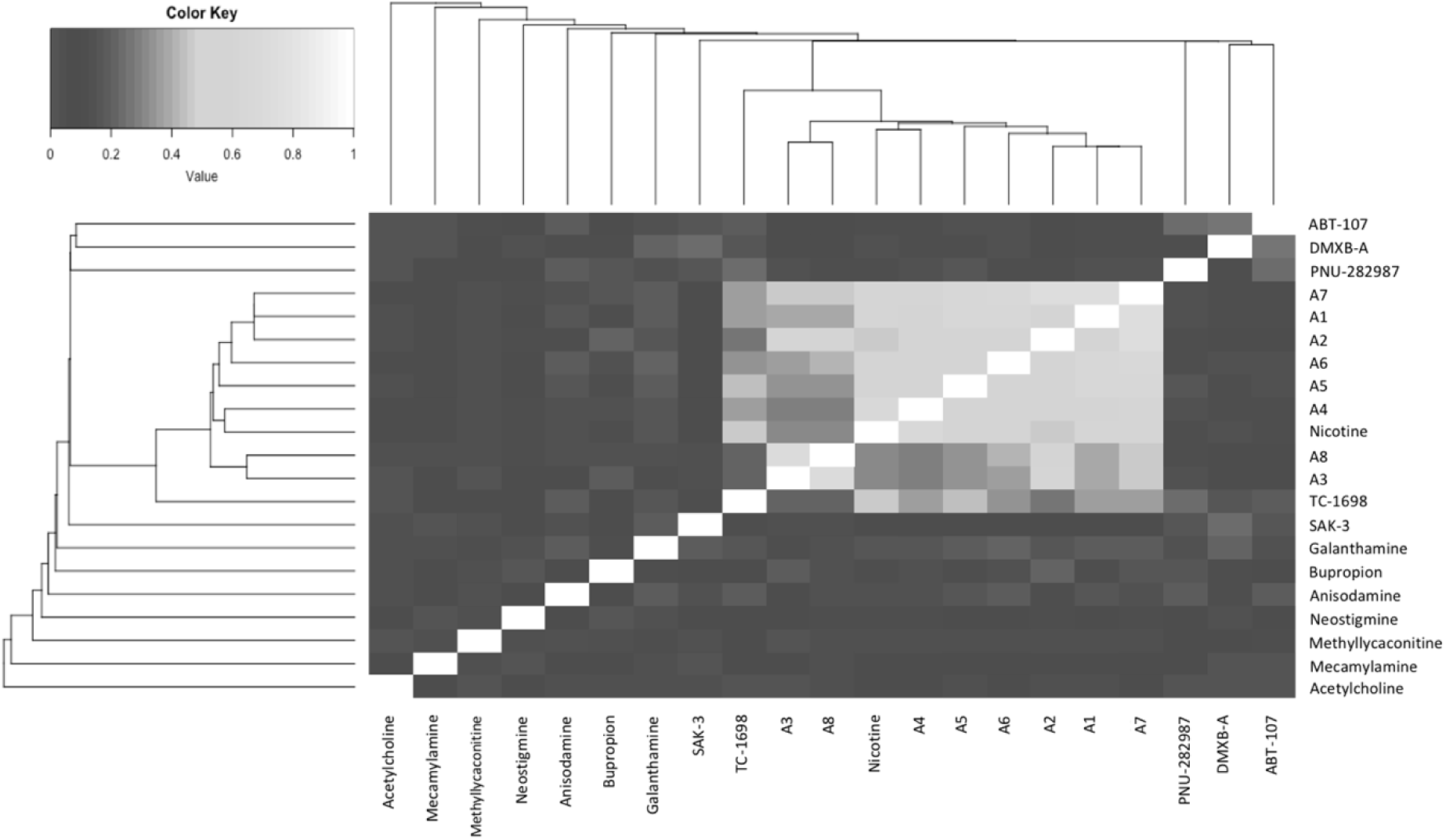
Similarity of molecules related to their potential neuroprotective response. The central cluster of the heatmap contains the analogs of nicotine, nicotine and TC-1698, both related to a neuroprotective *in vitro* positive activity over α7 nAChR.

**SUPPLEMENTARY Fig 2.**
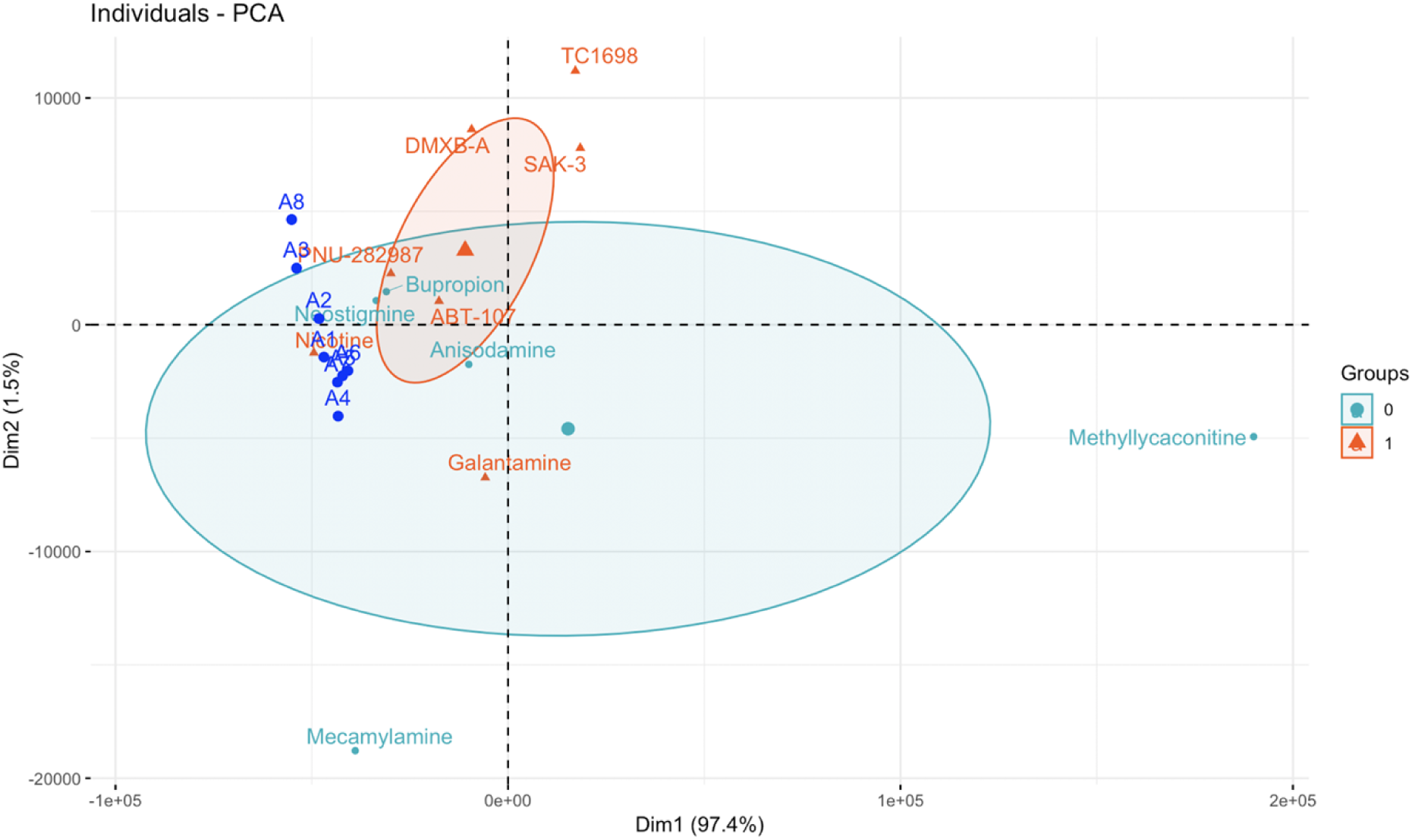
Principal component analysis of the manually curated dataset. 0 represents the antagonist function over the receptor (no neuroprotection) and 1 indicates the molecules related with a positive response of α7 nAChR and putative induction of PI3K/AKT Bcl-2.

**SUPPLEMENTARY Fig 3.**
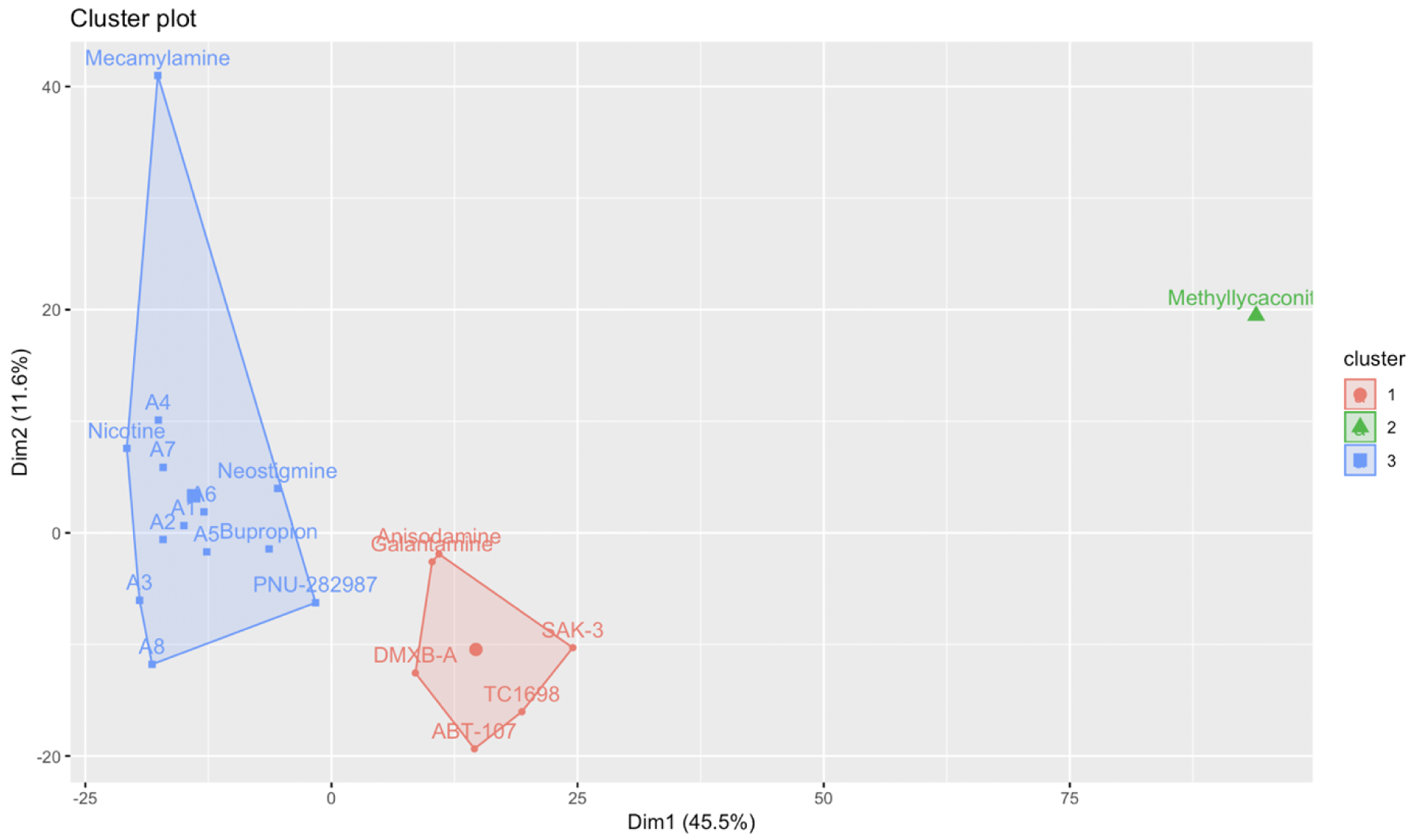
K-mean analysis of the manually curated dataset. The number of clusters were set to be self-organized and the clusters tend to organize the molecules by the activity over the receptor.

